# Exact solution of a three-stage model of stochastic gene expression including cell-cycle dynamics

**DOI:** 10.1101/2023.08.29.555255

**Authors:** Yiling Wang, Zhenhua Yu, Ramon Grima, Zhixing Cao

## Abstract

The classical three-stage model of stochastic gene expression predicts the statistics of single cell mRNA and protein number fluctuations as a function of the rates of promoter switching, transcription, translation, degradation and dilution. While this model is easily simulated, its analytical solution remains an unsolved problem. Here we modify this model to explicitly include cell-cycle dynamics and then derive an exact solution for the time-dependent joint distribution of mRNA and protein numbers. We show large differences between this model and the classical model which captures cell-cycle effects implicitly via effective first-order dilution reactions. In particular we find that the Fano factor of protein numbers calculated from a population snapshot measurement are underestimated by the classical model whereas the correlation between mRNA and protein can be either over-or underestimated, depending on the timescales of mRNA degradation and promoter switching relative to the mean cell-cycle duration time.

## I. INTRODUCTION

Over the past two decades, experiments have shown that gene expression is inherently noisy^1–6^ manifesting most visibly in the large variability of mRNA and protein numbers between cells. Understanding the origin of this noise and how its magnitude can be tuned by changes to the rate parameter values are important questions that can be addressed experimentally but also using mathematical modelling. Mathematical models of stochastic gene expression can be grouped into three classes: (i) those predicting only the statistics of mRNA numbers; (ii) those predicting only the statistics of protein numbers; (iii) those predicting both types of statistics including the correlation between mRNA and proteins. Of course the latter type is the most realistic and useful, however the other two are much more commonly found in the literature because of their simplicity and their use in interpreting data of only one of these two gene products, e.g. single-cell RNA sequencing^7,8^ and fluorescent labeling of proteins^9,10^. We next briefly review the large variety of models in each of these categories.

The simplest model of type (i) is the simple-birth death process whereby the birth reaction models transcription of mRNA and the death process models its subsequent decay via degradation and dilution. This is easily solved exactly and predicts a Poisson distribution in steadystate conditions^11^. However, a significant body of experiments suggests that the Fano factor (FF, the variance divided by the mean) for mRNA numbers is often substantially higher than that of a Poisson distribution^4,7,12^. To reconcile this discrepancy, a revision is necessary to the simple birth-death model which involves assuming that mRNA production occurs in bursts with a random size sampled from a geometric distribution; this can be solved exactly leading to a negative binomial distribution for mRNA numbers in steady-state conditions (which is well approximated by a gamma distribution when the mean mRNA numbers are not too small)^12–14^. However both of these models cannot explain the bimodality of distributions for mRNA numbers observed in some experiments^15^. A model that can explain bimodality and reduces to the birth-death or bursty birth-death models under certain conditions is the telegraph model^16,17^. This assumes that a promoter can switch between inactive and active states, and transcription occurs only from the latter; this model has been solved exactly in steady-state and in time, yielding an mRNA distribution in terms of hypergeometric functions. The commonest model of type (ii) is the bursty birth-death model which predicts steady-state negative binomial distributions for protein numbers (which is well approximated by a gamma distribution when the mean protein number is high)^18,19^. The main difference from the same model when used to predict mRNA distributions is the origin of bursting. For mRNA, bursts of expression occur when the promoter alternates between long transcriptional pauses and short intense periods of transcriptional activity. For proteins, bursts can arise from promoter switching and separately from the rapid translation of short-lived mRNA (transcriptional and translational bursting)^20^. Extensions of models of type (i) and (ii) to include an enhanced level of biological realism such as regular cell division events, gene dosage compensation, DNA replication, concentration homeostasis, cell-size homeostasis and cellto-cell variation in parameters, have also been developed and solved exactly or approximately under appropriate assumptions^13,21–30^.

The classical models of type (iii) are the two-stage and three-stage models of gene expression^20^. The twostage model has a simple birth-death process for mRNA production and degradation augmented with two additional first-order reactions describing protein translation and degradation/dilution. The joint distribution of mRNA and protein numbers of this model has been solved exactly in steady-state using the generating function method^31^; the time-dependent joint distribution has been obtained by invoking the partitioning property of Poisson processes^32^. A more realistic model is the three-stage model^3^ which is the two-stage model with the birthdeath model for mRNA noise replaced by the telegraph model. This model accounts for promoter switching, transcription, degradation and translation, and hence can be seen as the simplest model to capture the essential processes involved in the central dogma. While the moments of the gene products can be easily obtained^33^ (because the moment equations for any system of firstorder reactions are closed) there are no exact analytical solutions for this model’s marginal protein distributions and the joint distribution of mRNA and protein numbers (note that the marginal mRNA distribution is known and it is the same as that of the telegraph model). However even if these could be obtained, the issue remains that this model suffers from some obvious shortcomings, the most prominent of which is the fact that it does not explicitly model cell-cycle dynamics; for example it cannot capture the observation that typically in each cell, protein numbers increase between birth and cell division, followed by a sharp dip just after cell division^34^. This means that it is not clear how to relate the steady-state distribution predictions of the three-stage model to those from a population snapshot measurement, i.e. one that estimates the mRNA and protein in each cell at a point in time for a population of dividing cells with unsynchronized cell-cycles.

In this paper, we overcome these limitations. We extend the three-stage model to include a cell-cycle description and exactly solve for the time-dependent joint distribution of mRNA and protein numbers, thus providing a first dynamic stochastic description of both gene products within each cell-cycle and also across generations for two cases: a population of synchronized cells and the more typical population of unsynchronized cells where cell-age varies between cells. The population snapshot distribution of this model converges to a steady-state after a some generations; we numerically compare these to the steady-state solution from the classical three-stage model. In some regions of parameter space, we find large differences between the two models in their predictions of the protein Fano factor and of the correlation between mRNA and protein, thus highlighting the importance of an explicit cell-cycle description.

## II. RESULTS

### A. Models of stochastic gene expression

We consider three models of gene expression with increasing degrees of biological detail.

#### Model I

The classical three-stage model of stochastic gene expression (illustrated in the top left corner of Fig. 1) is described by the following reaction scheme

**FIG. 1.**
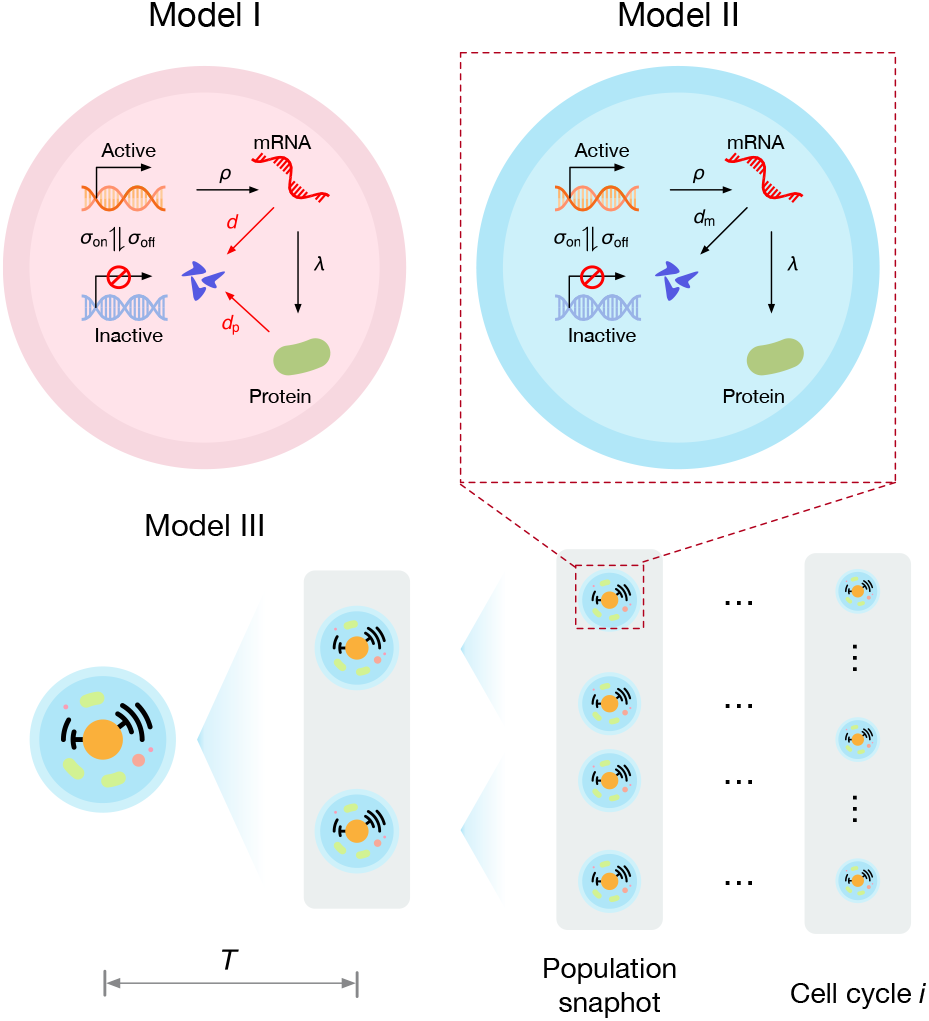
Illustration of three stochastic gene expression models. Model I: the classical three-stage model of stochastic gene expression that provides an effective description of promoter switching between active and inactive states, transcription, translation, mRNA decay and protein dilution. Model II: same as Model I except it does not describe protein dilution. This is a more realistic description of time-dependent gene expression within a cell-cycle than Model I since there is no protein dilution prior to cell division. Model III: this extends Model II by assuming that that a cell-cycle is of length *T* and that at cell division the mother cell’s mRNAs and proteins are randomly partitioned between its daughter cells. It provides a description of time-dependent gene expression within and across cell-cycles and is hence the most realistic model of the three.

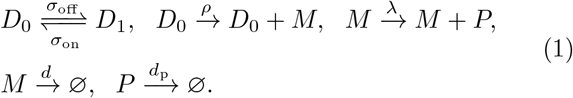

Here a promoter switches between two states: an active state *D*_0_ and an inactive state *D*_1_. The transitions between the two states occur at rates *σ*_off_ and *σ*_on_. Transcription occurs only from the state *D*_0_ leading to mRNA (*M*) production at a rate *ρ*. Proteins (*P*) are then translated from mRNA at a rate *λ*. mRNAs decay at a rate *d* (due to degradation and dilution), and protein dilution (due to cell division) takes place at a rate *d*_p_. Note that we cannot ignore active mRNA degradation because the lifetime of mRNA in mammalian cells is a fraction of the cell-cycle duration whereas for proteins the opposite is true: their lifetime due to degradation often exceeds the cell-cycle duration and hence degradation can be ignored in our model. For e.g. in mouse fibroblasts, the median lifetime of mRNA is 9 hours, the median lifetime of proteins is 46 hours and the cell-cycle duration is 27.5 hours^35^. All reactions in Eq. (1) are first-order and to be understood as being effective, i.e. describing the overall effect of thousands of elementary reactions enabling transcription, translation and decay processes inside a cell. The chemical master equation of this model has not been exactly solved; in particular an inspection of the generating function equations shows that the problematic term is that associated with the effective dilution reaction *P→* ∅.

#### Model II

In view of this issue we consider a simpler model (illustrated in the top right corner of Fig. 1) described by the following reaction scheme

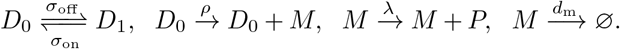

This is the same as Model I but without the protein dilution reaction; as well the mRNA decay rate here means specifically that due to degradation (has no contribution from dilution due to cell division) and hence we label it as *d*_*m*_ instead of *d*. As we shall see in the next section this is more mathematically tractable than Model I; however also it is a more biologically realistic model of time-dependent gene expression *within a single cell-cycle* (from birth to cell division) since there is no dilution prior to cell division.

#### Model III

Of course, generally we are not only interested in intra cell-cycle gene expression but also across many cell-cycles. To describe this case we extend Model II by assuming that at the end of the cell-cycle, mRNA and protein molecules are randomly partitioned with probability 1*/*2 between two daughter cells. Specifically, just after cell division, the probability of a daughter cell having *n*_m_ mRNAs and *n*_p_ proteins, conditional on the mother cell having 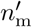 mRNAs and 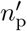 proteins, is given by

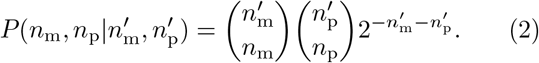

While being a simple model of the complex processes occurring at cell division, this is clearly far more realistic than the first-order protein dilution reaction in Model I since dilution now occurs at a point in time rather than continuously. Model III is illustrated at the bottom of Fig. 1.

The probabilistic description of Model II is given by the CME

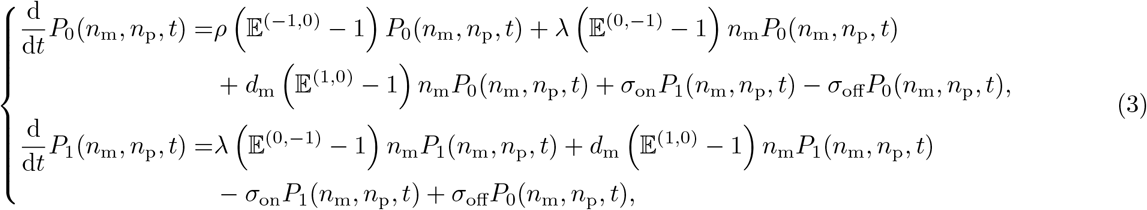

where *P*_*φ*_(*n*_m_, *n*_p_, *t*) is the probability of observing *n*_m_ mRNAs and *n*_p_ proteins in a cell when a promoter is active (*φ* = 0) or inactive (*φ* = 1). The step operator 𝔼 ^(*i,j*)^ acts on a general function *f* (*n*_1_, *n*_2_) as 𝔼 ^(*i,j*)^*f* (*n*_1_, *n*_2_) = *f* (*n*_1_ + *i, n*_2_ + *j*)^36^. For compactness of notation, the time argument *t* is hereafter omitted. Next, we show how to solve Eq. (3) by means of the generating function method.

By defining the generating function 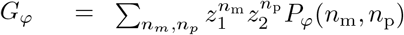 Eq. (3) can be recast as the following set of partial differential equations

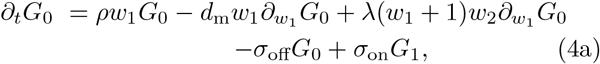

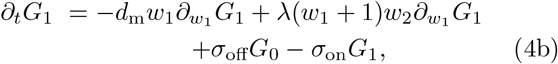

where *w*_1_ = *z*_1_− 1 and *w*_2_ = *z*_2_− 1. For ease of notation, the arguments of the generating function *w*_1_ and *w*_2_ are omitted. Once the generating function equations are solved, the conditional joint distributions *P*_*φ*_(*n*_m_, *n*_p_) in Eq. (3) can be constructed using the relation

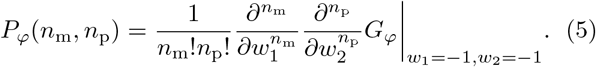

We note that because the evaluation of high-order symbolic derivatives can be troublesome, a more practical approach is needed. We first symbolically expand the generating function as a Taylor series of *w*_1_ using the Mathematica command Series and then perform a Taylor series expansion to the 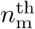 coefficient for *w*_2_ using the Mathematica command NSeries. The probability *P*_*φ*_(*n*_m_, *n*_p_) is then simply given by the 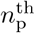 coefficient. A similar approach has previously been used for the rapid evaluation of marginal distributions from the generating function^37,38^.

We start by using Eq. (4a) to write *G*_1_ as a function of *G*_0_:

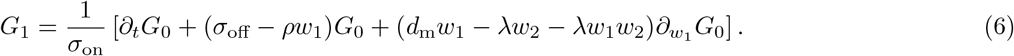

Summing up Eqs. (4a) and (4b), and plugging Eq. (6) into the resultant equation, we have a parabolic partial differential equation that depends only on *G*_0_:

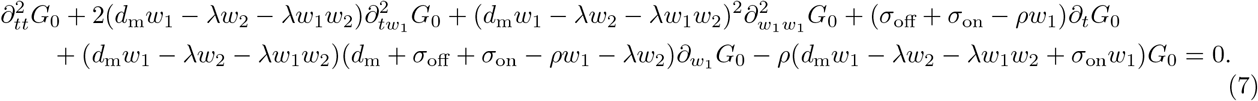

Note that there is no partial derivative with respect to *w*_2_ in Eq. (7), which suggests that the solutions to Eq. (7) live in a space spanned by the coordinates (*t, w*_1_). To solve Eq. (7), we convert the coordinates (*t, w*_1_) to a new set of coordinates (*u, v*) as

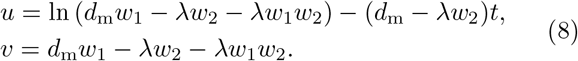

This is a standard coordinate-conversion technique for solving parabolic PDEs^39^. The following relations between the derivatives of the two sets of coordinates hold

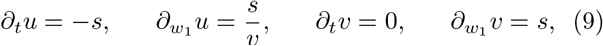

where *s* = *d*_m_ *λw*_2_.. We use the chain rule and Eq. to convert the firstand second-order derivatives of *G*_0_ under the coordinates (*t, w*_1_) to those under the new coordinates:

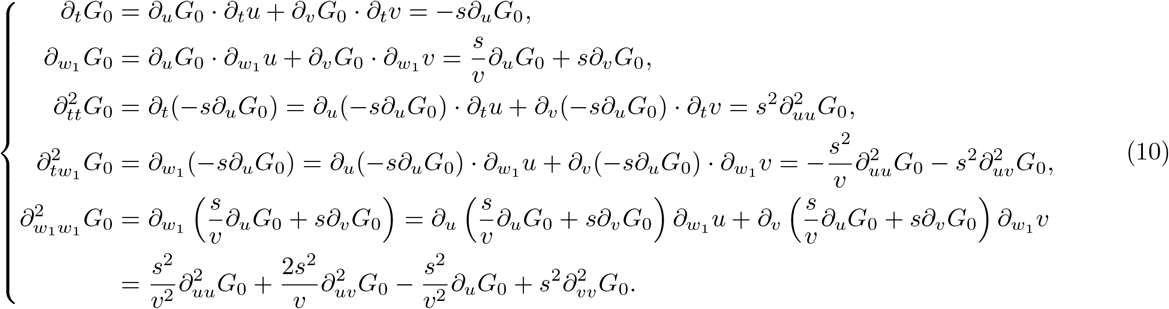

By using these equations, the parabolic PDE Eq. (7) further reduces to a second-order ODE

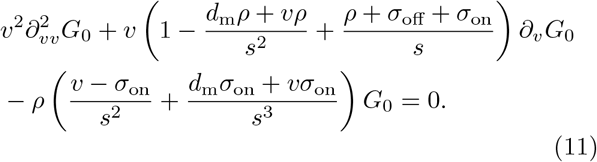

Note the coefficient of *G*_0_ in Eq. (11) is *v*-dependent, which prevents us formulating the solution as standard solutions of some canonical second-order ODEs. To circumvent this issue, we replace *G*_0_ in Eq. (11) with

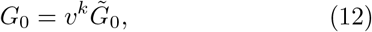

which simplifies Eq. (11) to

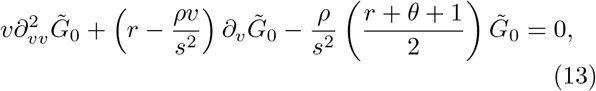

where *r, θ* and *k* are defined as

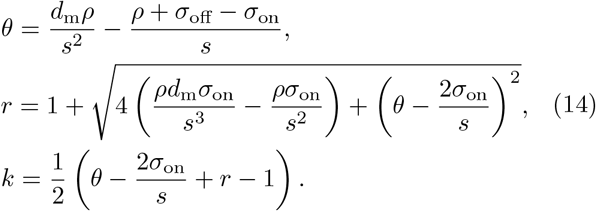

Changing to a description in terms of the variable *x* = *ρv/s*^2^, Eq. (13) further simplifies to

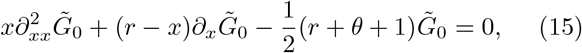

which is the canonical form of Kummer’s equation (see Eq. (13.2.1) in Ref. 40). This suggests the general solution to Eq. (15) is

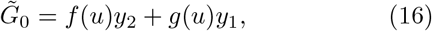

where *y*_1_ and *y*_2_ have the form:

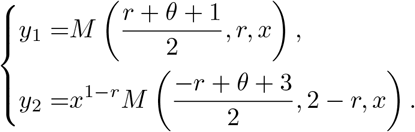

Note that *M* is the confluent hypergeometric function, also known as Kummer’s function, and the functions *f* and *g* only depend on *u*, and are to be determined from the initial conditions. It immediately follows from Eqs. (12) and (16) that the general form of *G*_0_ is

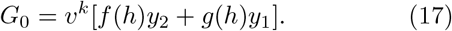

Note that the functions *f* and *g* depend on *u* in Eq. (16), while they depend on *h* = exp(*u*) in Eq. (17). Since *h* is a function of *u* only and *f* and *g* are undetermined, we slightly abuse the *f* -*g* notations in Eq. (17) to facilitate the following calculations.

Next we solve for *G*_1_ as a function of *f, g, y*_1_ and *y*_2_. This can be done via Eq. (6), and it requires the calculation of ∂_*t*_*G*_0_ and ∂_*w*_ *G*_0_. Combined with *u* = ln *v*− *st* and *h* = *v* exp(−*st*), the two derivatives can be calculated according to Eq. (10) as follows

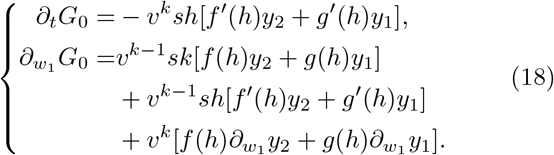

Plugging Eqs. (17) and (18) into Eq. (6), we find

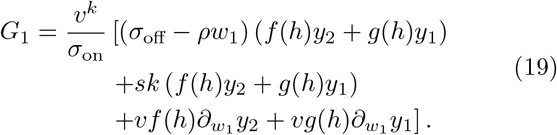

By using one of the derivative relations of Kummer’s function, i.e. Eq. (13.3.15) in Ref. 40, the terms ∂_*w1*_ *y*_1_ and ∂_*w1*_ *y*_2_ in Eq. (19) can be expressed in a form that is free of derivatives over Kummer’s function, which easies their numerical evaluation:

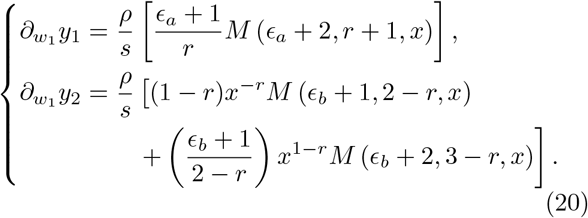

Here the two quantities *ϵ*_*a*_ and *ϵ*_*b*_ are defined as

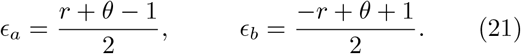

Hence it can be shown (Appendix A) that Eq. (19) simplifies to

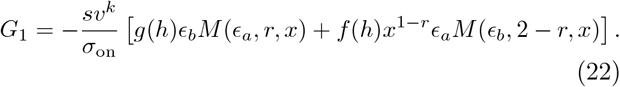

All that remains to obtain a full solution to Model II is to determine *f* and *g* in Eqs. (17) and (22). We assume that the initial distribution solutions of the CME Eq. (3), conditioned on the promoter state, are given by *p*_0_(*n*_m_, *n*_p_) and *p*_1_(*n*_m_, *n*_p_). One can immediately deduce that the corresponding initial conditions for Eqs. (4a)-(4b) are given by

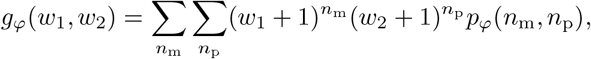

for *φ* = 0 or 1. At time *t* = 0, the following relation also holds

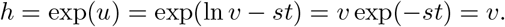

Then, using Eqs. (17) and (19) one can establish the link between the initial conditions *g*_0_(*w*_1_, *w*_2_), *g*_1_(*w*_1_, *w*_2_), and the functions *f* and *g*:

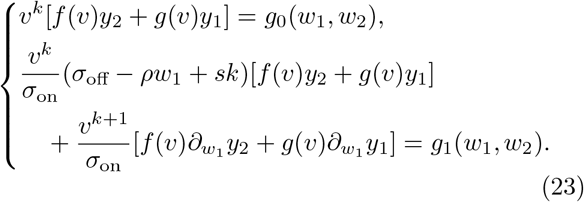

Solving these equations simultaneously leads us to

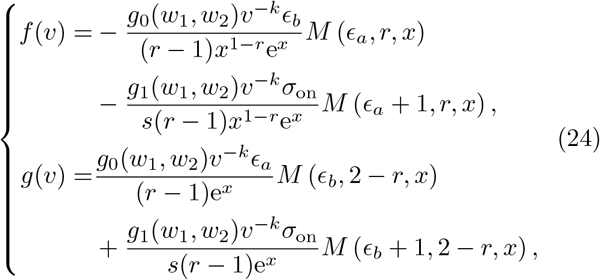

with the condition *h* = *v* when *t* = 0 (doe detailed steps of the calculation see Appendix B).

Now, *f* and *g* in Eq. (24) are functions of the variable *h* (see Eq. (17)); it is also the case that *h* = *v* exp(*st*) and *h* = *v* at time *t* = 0. Hence to obtain the solution of *f* and *g* for any time *t*, we have to replace *v* by *h*, i.e., *v→ v* exp(− *st*) and *x→ x* exp(− *st*). Since the coordinate *w*_1_ is dependent on *v* (*w*_2_ is not a coordinate), we also need to seek a similar mapping 𝒯_*t*_ for *w*_1_. From Eq. (8), it is known that

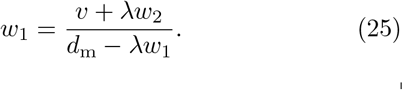

Further applying the mapping *v 1→ v* exp(−*st*), we obtain the mapping *𝒯*_*t*_

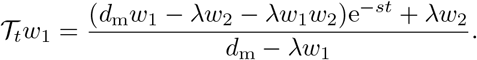

Applying the three mappings *v → v* exp(−*st*), *x → x* exp(−*st*) and *w*_1_ *→ 𝒯*_*t*_*w*_1_ to Eq. (25), and combining with Eqs. (17) and (A1), we finally find the full exact time-dependent solution to Model II, which is summarized below as Eqs. (26) and (28).

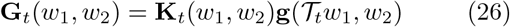

where **G**_*t*_(*w*_1_, *w*_2_) = [*G*_0_(*w*_1_, *w*_2_, *t*), *G*_1_(*w*_1_, *w*_2_, *t*)]^⊤^, **g**(*𝒯*_*t*_*w*_1_, *w*_2_) = [*g*_0_(*𝒯*_*t*_*w*_1_, *w*_2_), *g*_1_(*𝒯*_*t*_*w*_1_, *w*_2_)]^⊤^, and

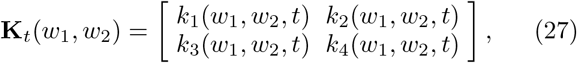

with

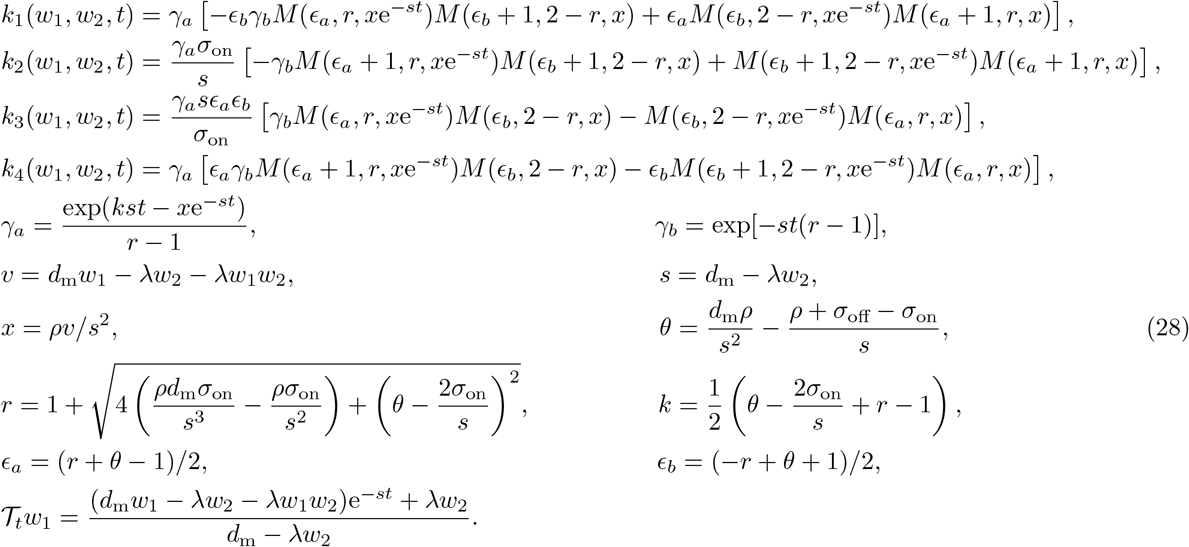

As mentioned earlier, the joint distribution of mRNA and protein numbers follow directly from Eq. (5).

#### Special Case (i)

Consider the system initiated without any mRNA and protein and in the active promoter state. The generating functions for these initial conditions are *g*_0_(*w*_1_, *w*_2_) = 0 and *g*_1_(*w*_1_, *w*_2_) = 1. The general solution given by Eqs. (26) and (28) then greatly simplifies to

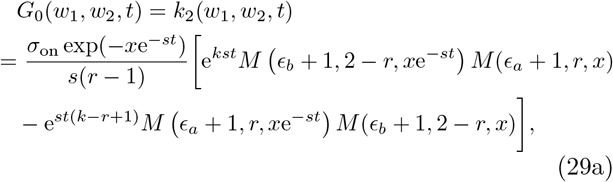

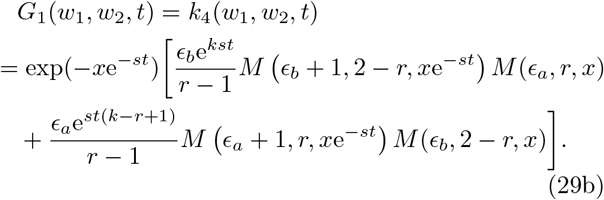

#### Special Case (ii)

The marginal distribution of mRNA numbers conditioned on the promoter state can be obtained by setting *w*_2_ to 0 in Eqs. (26) and (28); as expected, this is exactly equal to the solution of the the well known telegraph model^17^ since the protein production and decay reactions in Model II have no influence on mRNA dynamics (for detailed calculations see Appendix C).

#### Special Case (iii)

Similarly setting *w*_1_ to 0 gives us the marginal protein number distribution conditioned on the promoter state which to the best of our knowledge has not been derived before. This distribution considerably simplifies when promoter-inactivation events occur far more frequently than promoter-activation events, i.e., *σ*_off_ ≫ *σ*_on_; and that mRNA decays fast, i.e. *d*_m_ ≫ 0. This is indeed the exact solution to the chemical master equation of the reduced reaction system

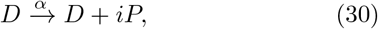

where *i* is a random number sampled from the geometric distribution *P* (*β, i*) = *β*^*i*^*/*(1 + *β*)^*i*+1^ where *i* = 0, 1, 2, … Note that *β* is the mean burst size. The kinetic parameters are given by

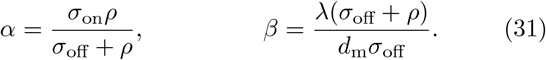

For details of the derivation see Appendix D.

#### Numerical validation of the exact solution

In Figs. 2 and 5, we show that the exact analytical solutions for the joint and marginal distributions are in excellent agreement with those obtained using the stochastic simulation algorithm (SSA)^41^ when there is no bimodality (top row) and when there is bimodality (bottom row).

**FIG. 2.**
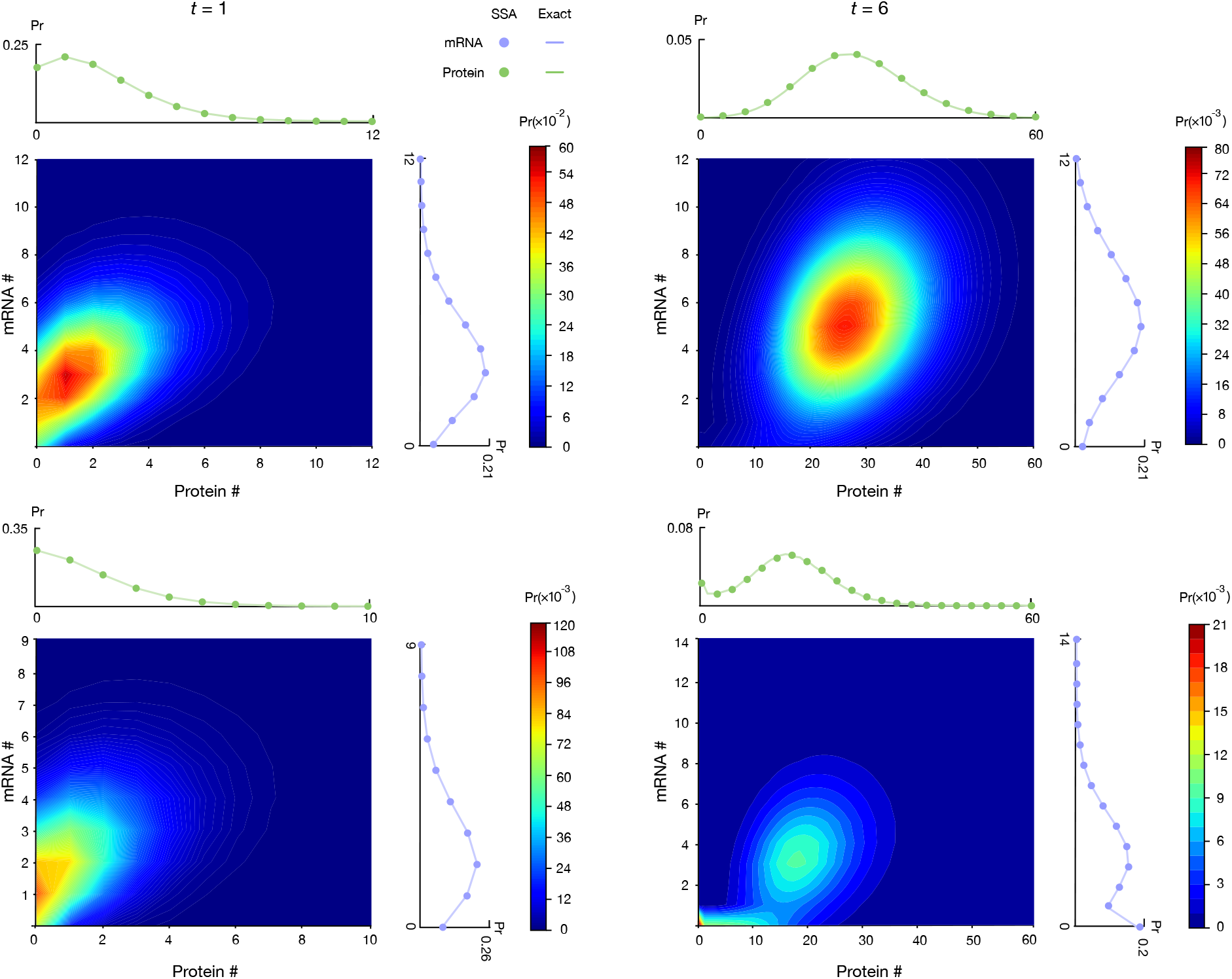
Comparing the analytical joint and marginal distributions of mRNA and protein numbers in Model II predicted by Eqs. (26) and (28) with those obtained from stochastic simulations using the SSA (the number of independent realizations is 10^7^). The heatmaps show the joint distributions of mRNA and protein numbers predicted by the analytical solution at time *t* = 1 and *t* = 6, and they agree well with the predictions given by SSA in Fig. 5. The marginal distributions of mRNA numbers given by the exact solution (solid lines) perfectly match the ones given by the SSA (dots). Similar results are found for the marginal distributions of protein numbers. The kinetic parameters are (top panel) *ρ* = 6, *λ* = 1, *σ*_off_ = 0.1, *σ*_on_ = 1, and *d*_m_ = 1; (bottom panel) *ρ* = 4, *λ* = 1, *σ*_off_ = 0.05, *σ*_on_ = 0.02, and *d*_m_ = 1. Initially the promoter is in the ON state and there are zero mRNA and protein numbers.

## III. EXACT SOLUTION FOR THE TIME-DEPENDENT JOINT DISTRIBUTION OF MRNA AND PROTEIN NUMBERS OF MODEL III

Next we solve Model III which is an extension of Model II. In particular in Model III we introduce a cell-cycle description whereby at the end of each cell-cycle the gene products are randomly partitioned between daughter cells. For generality, we shall assume that the duration of the *n*^th^ cell-cycle is *T*_*n*_.

We start by setting the initial conditions of Model II, *g*_0_ and *g*_1_, to be the two generating functions at the beginning of cell-cycle *n*. Hence using Eq. (26) we have the time-evolution equations of the two generating functions within cell-cycle *n* (prior to cell division)

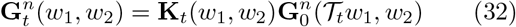

where 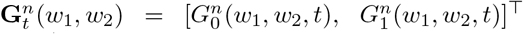, 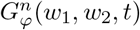 is the generating function of promoter state *D*_*φ*_ at cell age *t* ∈ [0, *T*_*n*_) of cell-cycle *n*, and The next step is to seek a recursive equation between 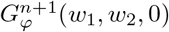 and 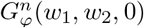. According to the definition of the generating function, we have

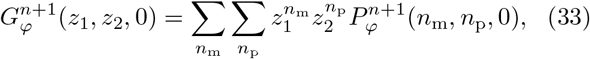

where we temporarily use *z*_1_ and *z*_2_ instead of *w*_1_ and *w*_2_ to facilitate the following derivation. The probabilities before and after cell division are linked by the law of total probability

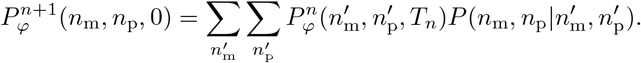

Using the binomial partitioning condition Eq. (2), Eq. (33) can be written as

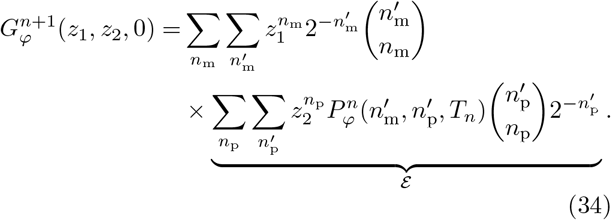

The term *ε* can be further simplified to

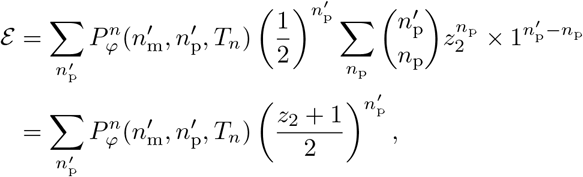

where the second step follows from the binomial theorem. By performing the same simplification steps Eq. (34) reduces to

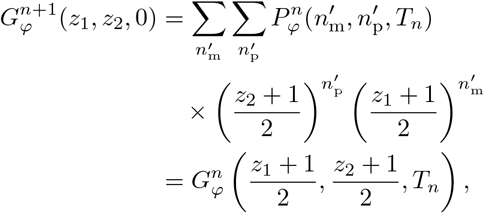

where the last step uses the definition of the generating function. Switching *z*_1_ and *z*_2_ back to *w*_1_ and *w*_2_, we finally obtain the recursive equation linking the generating function prior to cell division to that after cell division

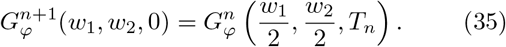

In summary, taken together, Eqs. (27), (32) and (35) constitute the exact solution to Model III. We note that in this description, at all points in time, only one cell is tracked, i.e. when a cell divides, only the gene product contents of one of the daughter cells is followed. Hence each independent realisation of the SSA simulating Model III describes a particular lineage; this means that the total number of cells in the population does not change with time. If furthermore *T*_*n*_ = *T*, the population of cells has perfectly synchronized cell-cycles. In Fig. 3, we perform 4 sets of simulation experiments. In each experiment, for simplicity we choose *T*_*n*_ = *T*, i.e. the cellcycle duration is the same for all cycles; *T* is however different for each of the four experiments. We compare the marginal distribution of protein counts predicted by our exact solution of Model III and the SSA at different cell ages in the first 4 cell-cycles. As expected, the theoretical predictions and the SSA are found to be in excellent agreement verifying that our theory accurately predicts gene expression within and across cell-cycles. Note also that here we have focused on testing the accuracy of the protein number distribution because the mRNA number distribution for Model III s a special case of an extended version of the telegraph model studied earlier^21^. The theoretical joint distribution of mRNA and protein numbers of Model III can also be directly compared with the SSA and again the two are found to be in excellent agreement (results not shown).

**FIG. 3.**
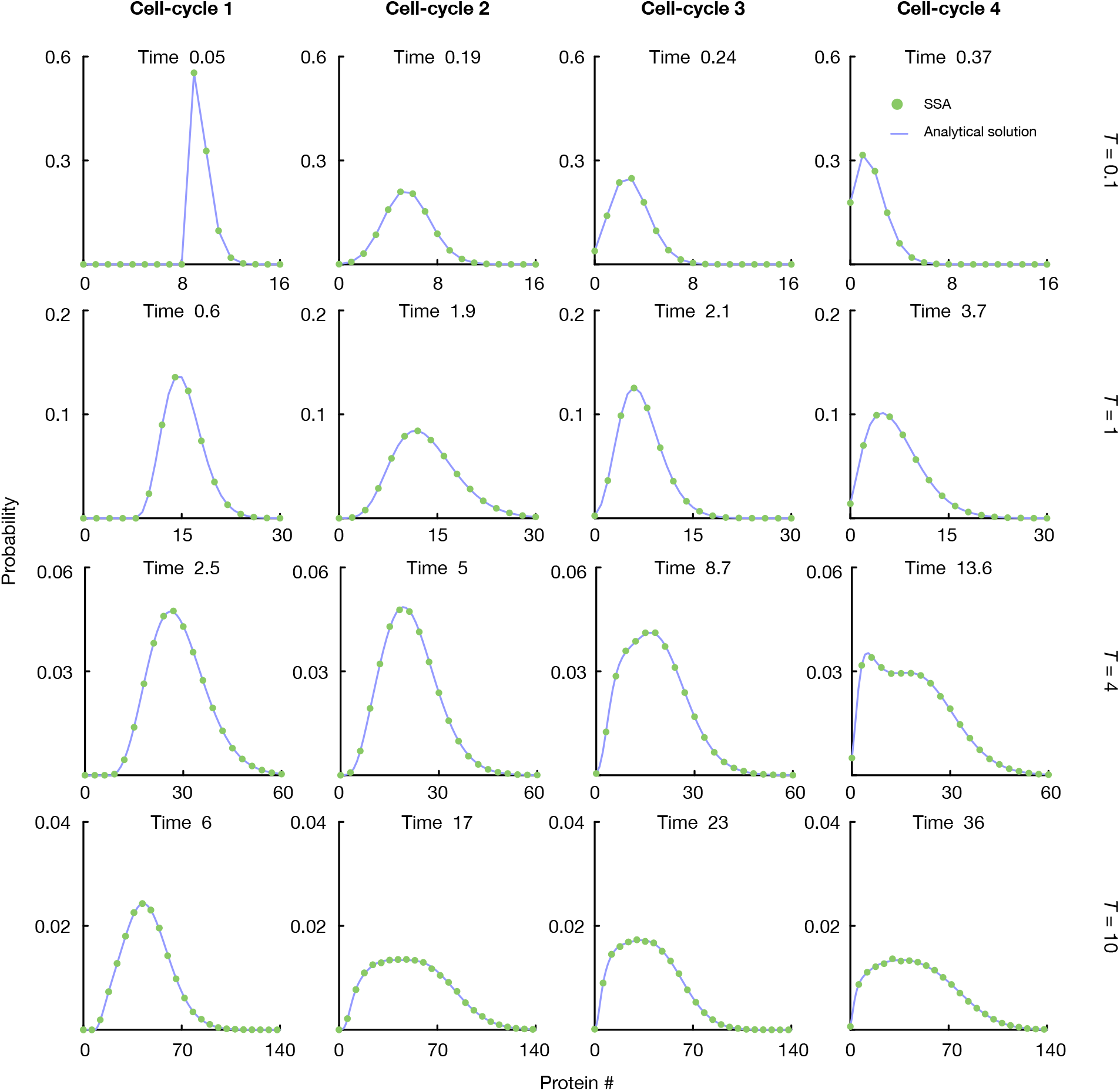
The marginal distributions of protein numbers in Model III given by theory (Eqs. (27), (32) and (35)) are in excellent agreement with the distributions computed using the SSA at different cell ages and for different cell-cycle duration *T*. The time shown in the figures is the absolute time measured from the beginning of the first cell-cycle. In all the cases, the system is initiated in the active promoter state *D*0 with 4 mRNA and 9 protein molecules. For simplicity, the cell-cycle duration is chosen to be the same for all cell-cycles, i.e. *T*_*n*_ = *T*. The kinetic parameters are *ρ* = 2, *λ* = 3, *σ*_off_ = 0.1, *σ*_on_ = 0.1 and *d*m = 1. We use the modified version of the SSA in Ref. 13, and the number of independent realizations is 10^6^.

## IV. COMPARISON OF THE STATIONARY GENE PRODUCT NUMBER STATISTICS OF MODELS I & III

### Model I

The steady-state statistics of Model I are easy to obtain directly from the moment equations derived from the CME^33^. This is because the moment equations for any system of first-order reactions are closed and hence their solution in steady-state simply amounts to solving a system of simultaneous equations^42^. The means, variances, Fano factors (ratio of variance to mean of the molecule numbers), the covariance of mRNA and protein number fluctuations, and the correlation coefficient are given by:

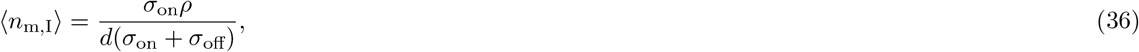

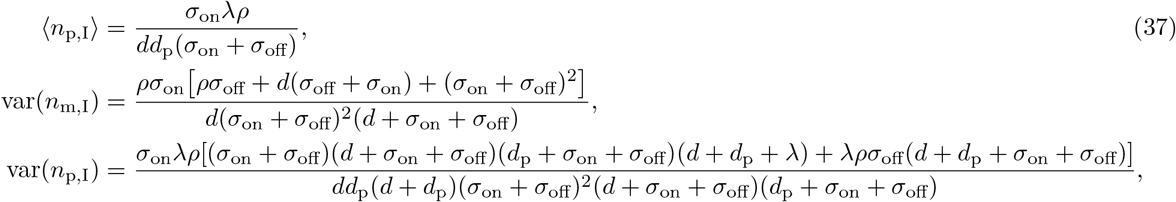

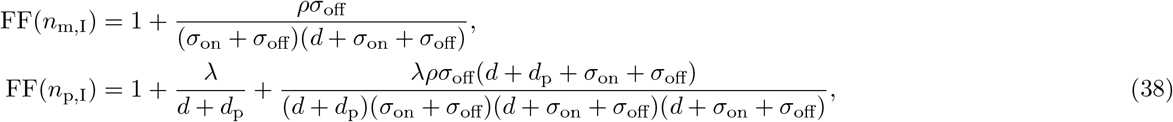

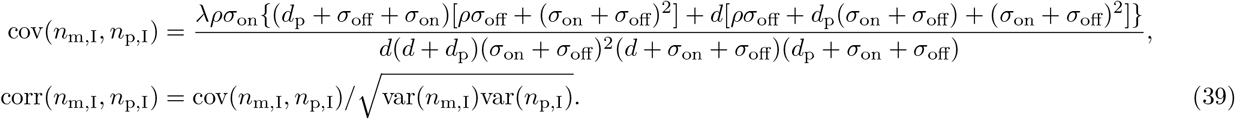

### Model III

To compare with the steady-state statistics of Model I, we first need to define what is precisely meant by the steady-state of Model III. We shall consider the common case where measurements of the mRNA and protein numbers for each cell are done at a point in time for a (growing) population of dividing cells with unsynchronized cell-cycles. Furthermore these measurements are done when steady-state growth has been achieved, i.e. the probability that a cell of age *t* has a given number of mRNA and proteins is independent of which generation it belongs to^43^.

The calculations leading to the derivation of the distributions of mRNA and proteins for a population snapshot measurement are complex and will be done in two steps: (i) first we will enforce steady-state growth conditions on Model III, a condition also referred to as the cyclostationary condition in the literature^26^. This will lead to a steady-state distribution of gene products for cells of age *t* where *t∈* [0, *T*]; (ii) the distributions obtained from the former step will then be integrated over the cellage distribution in a population thus leading to the final result.

#### 1. Enforcing steady-state growth

According to Eqs. (32) and (35), the generating functions at cell age *t* = 0 of two successive cell-cycle are related by

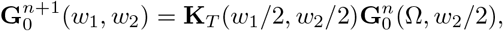

with

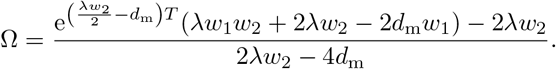

Then, the cyclo-stationary condition suggests that there exists a limit such that 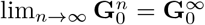. It is equivalent to 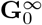 satisfying

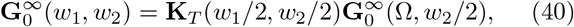

which means the probability distribution of mRNA and protein numbers at cell age *t* = 0 reaches steady state. Once the cyclo-stationary condition is satisfied for *t* = 0, the generating functions 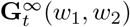 also becomes cyclostationary, since **K**_*t*_ is cell-cycle invariant in

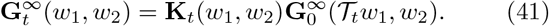

By taking derivative of *w*_1_ on the both sides of Eq. (40) and setting *w*_1_ = *w*_2_ = 0, one can get the equations governing the cyclo-stationary means of mRNA numbers at cell age *t* = 0

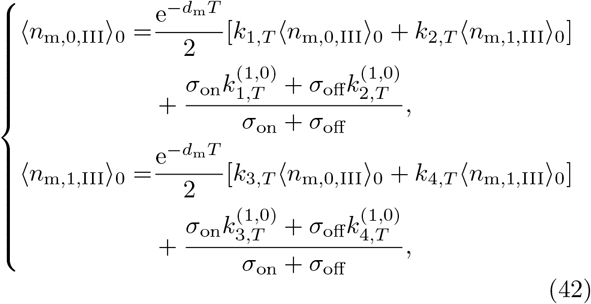

where ⟨ *n*_m,*φ*,III_ ⟩ _*t*_ is the cyclo-stationary mean mRNA number at promoter state *D*_*φ*_ and cell age *t*, and we use the following short-hand notations

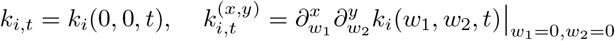

for *i* = 1, 2, 3, 4. The total mean mRNA number at cell age *t, ⟨n*_m,III_*⟩*_*t*_, is then equal to

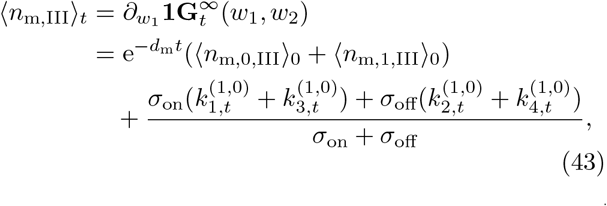

where **1** = [1, 1]. Solving for ⟨ *n*_m,0,III ⟩ 0_ and ⟨*n*_m,1,III_ ⟩_0_ from Eq. (42) and substituting them into Eq. (43), we find an analytical expression for the mean mRNA number at cell age *t* under cyclo-stationary conditions

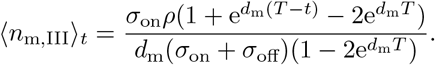

By taking derivative of *w*_2_ on the both sides of Eqs. (40) and (41) and using the same principles, we can also get the expression for protein (⟨*n*_p,III_ ⟩_*t*_)

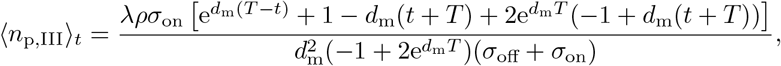

Similarly one can find the second-order raw moments for mRNA numbers and protein numbers under the cyclo-stationary condition 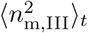, 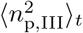 and 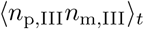. These are too complex to write here (the interested reader is referred to the Mathematica notebook).

#### 2. Taking into account the lack of cell-cycle synchronization between cells

Finally we need to take into account the fact that often the cell-cycles of different cells are not synchronized and hence a population snapshot will measure gene product numbers from cells of different ages. The issue is that if we consider stochastic simulations of Model III then if we start from a single cell, at any one time in the future we obtain a population of perfectly synchronized cells due to the deterministic cell-cycle duration. Of course, in reality cell-cycle duration is not deterministic but rather is a random variable, a property that will naturally desynchronize cells. It can be shown that if the cell-cycle duration has some small noise about a mean *T* then the cellage distribution is given by *p*(*t*) = 2^1−*t/T*^ ln 2*/T* where *t ∈* [0, *T*]^27,43^. Note that this implies that it is more probable to observe cells of a young age (a direct implication of the doubling of the number of cells with age *t* = 0 when cell division occurs).

Hence the observed mean mRNA numbers for a cell population (⟨*n*_m,III_⟩) is given by taking the ensemble average of the cyclo-stationary mean over *p*(*t*). This leads to

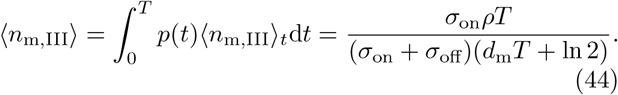

The stationary mean protein number for the cell population can be similarly computed

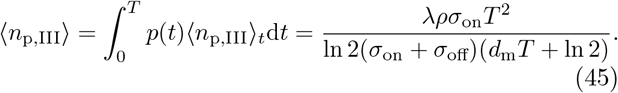

The variance of mRNA and protein numbers for the cell population are similarly given by

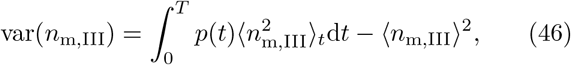

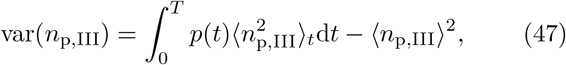

while the covariance of mRNA and protein fluctuations is given by

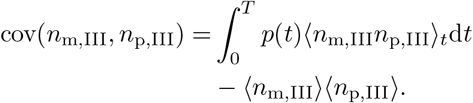

The correlation between mRNA and protein is then obtained using

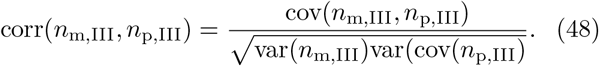

The expressions derived above were validated using stochastic simulations, as follows. As mentioned earlier, a non delta-function cell-age distribution can only arise if the cell-cycle duration is not fixed. To this end, we devise Model IV, one that is the same as Model III except that the cell-cycle duration is equal to the mean *T* plus small noise which is Erlang distributed (an assumption that is motivated by measurements of several cell types; see Fig.1c in Ref. 27). For a more detailed discussion of Model IV we refer the reader to Appendix E. Simulations of Model IV leads to an exponentially increasing population of cells; at a point in time after several generations have elapsed, we calculate the means, variances and covariance of mRNA and proteins across the cell population and also the cell age distribution. In Fig. 6 these are compared with those predicted by Model III; excellent agreement is found for three parameter sets thus validating the accuracy of Eqs. (44)-(48).

### Comparison of Models I and III

Now we are in a position to compare Models I and III so that we can assess the importance of modelling the cell-cycle explicitly. To ensure a fair model comparison, we first equate the means of the two models, i.e. ⟨*n*_m,I_⟩=⟨*n*_m,III_⟩ and ⟨*n*_p,I_⟩ = ⟨*n*_p,III_ ⟩. This equivalence is possible provided the effective mRNA decay rate *d* and the effective protein dilution rate *d*_p_ in Model I, are chosen to be

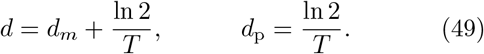

Using these effective rates, we compute the Fano factor for proteins in Model I, FF(*n*_p,I_), using Eq. (38). The Fano factor for proteins in Model III, FF(*n*_p,III_), can be computed from Eqs. (45) and (47). We then introduce the ratio *R*_1_ defined by

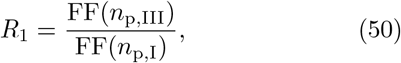

which quantifies the discrepancy between the two Fano factor predictions provided by Model I and Model III. Similarly, we define the ratio *R*_2_ as

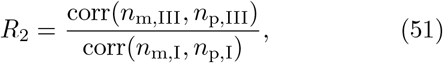

to quantify the discrepancy between the correlation coefficients between mRNA and protein numbers predicted by Model I (Eq. (39)) and Model III (Eq. (48)).

In Fig. 4a and b, we investigate how the ratio *R*_1_ varies across parameter space. The results show that Model I always underestimates the Fano factor of protein number fluctuations. The deviations between the two models is mostly determined by the ratio of the time spent in the ON and OFF states and the cell-cycle duration time (*σ*_off_*T* and *σ*_on_*T*, respectively) and to a much lesser extent by the ratio of the mRNA lifetime and the cell-cycle duration time *d*_*m*_*T*. Generally, the deviations increase with *σ*_on_*T* and with decreasing *σ*_off_*T* and *d*_*m*_*T*. In Fig. 4c and d, we investigate how the ratio *R*_2_ varies across parameter space. The results show that the Model I can either overestimate or underestimate the correlation coefficient. The factor that most strongly determines whether *R*_2_ is less than or greater than 1 is the ratio of the mRNA lifetime and the cell-cycle duration time *d*_*m*_*T*; we note that the line *d*_*m*_*T* = 10 is approximately coincident with *R*_2_ = 1.

**FIG. 4.**
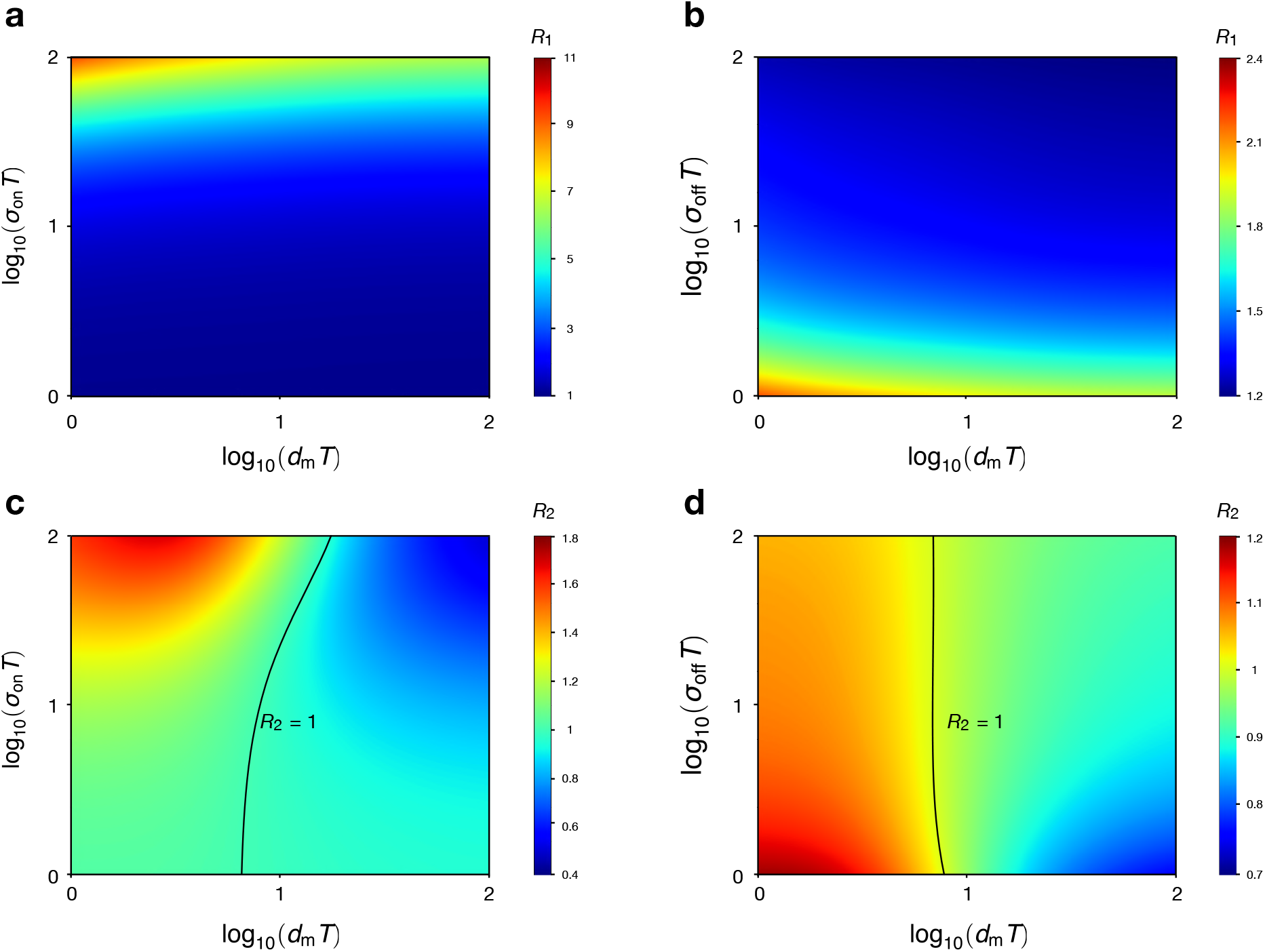
Comparison of the steady-state Fano factor of protein numbers and of the correlation coefficient between mRNA and protein predicted by Model I and Model III. The ratio *R*1 is the Fano factor of Model III divided by that of Model I (Eq. (50)). The ratio *R*2 is the correlation coefficient of Model III divided by that of Model I (Eq. (51)). In (a) and (c) we fix the values of the switching off rate *σ*_off_ = 1.5 hr^−1^ and the transcription rate *ρ* = 7 hr^−1^ to be the medians reported in Ref. 7, the translation rate *λ* = 198 hr^−1^ mRNA^−1^ to be the median reported in Ref. 35, and the cell-cycle duration *T* = 27.5 hrs to be the value reported in Ref. 35. In (b) and (d) the parameter values are the same, except that now we fix the switching on rate *σ*_off_ = 0.12 hr^−1^ to be the median reported in Ref. 7. Note that *R*_1_ *>* 1 all over parameter space implying that Model I always underestimates the Fano factor of protein fluctuations. The ratio *R*_2_ can be greater than or less than 1 implying that Model I can either over or underestimate the correlation coefficient between mRNA and protein numbers.

**FIG. 5.**
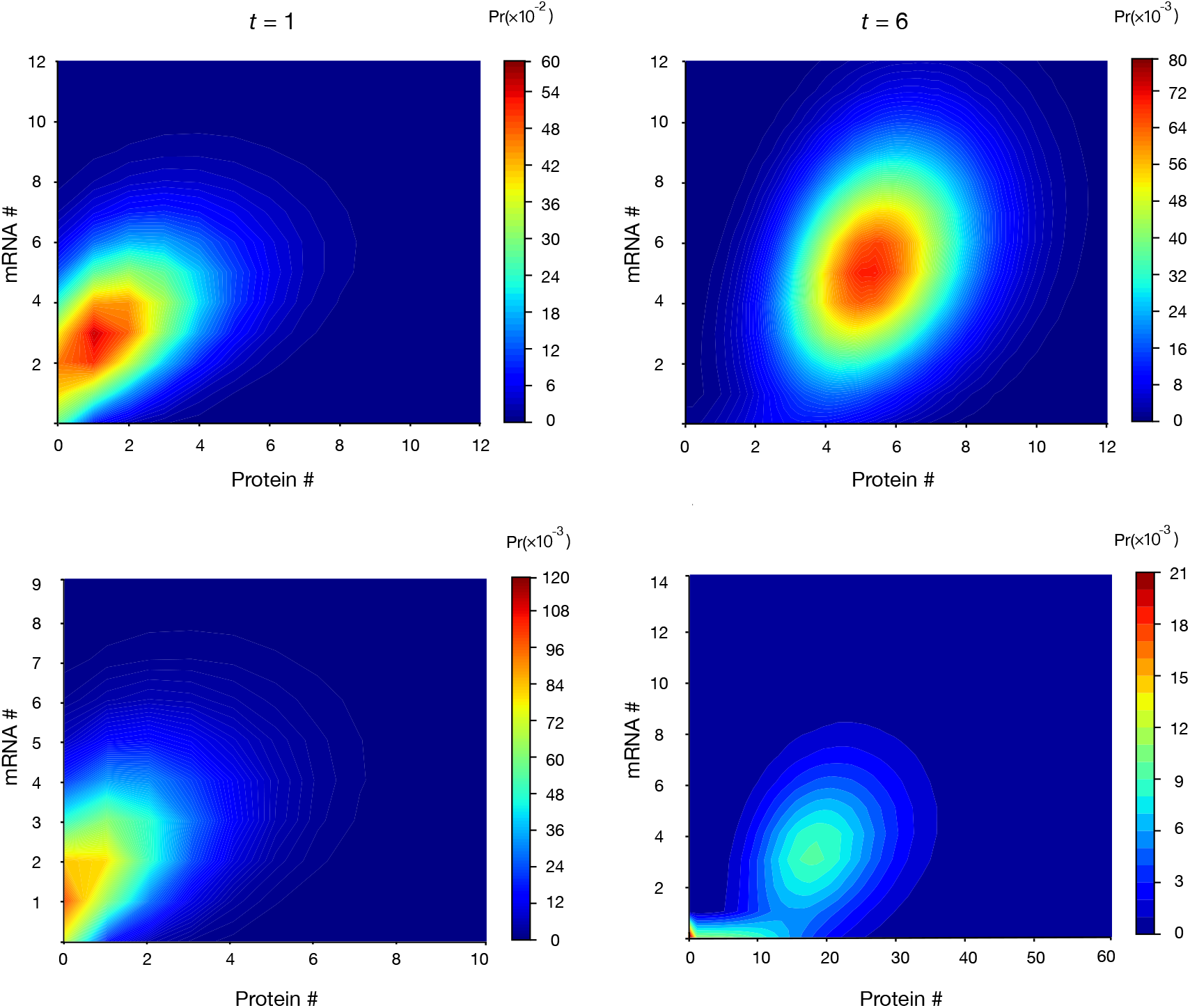
Joint distributions of mRNA and protein numbers in Model II, as predicted by the SSA (10^7^ realizations). The kinetic parameters and initial conditions are the same as those in Fig. 2.

**FIG. 6.**
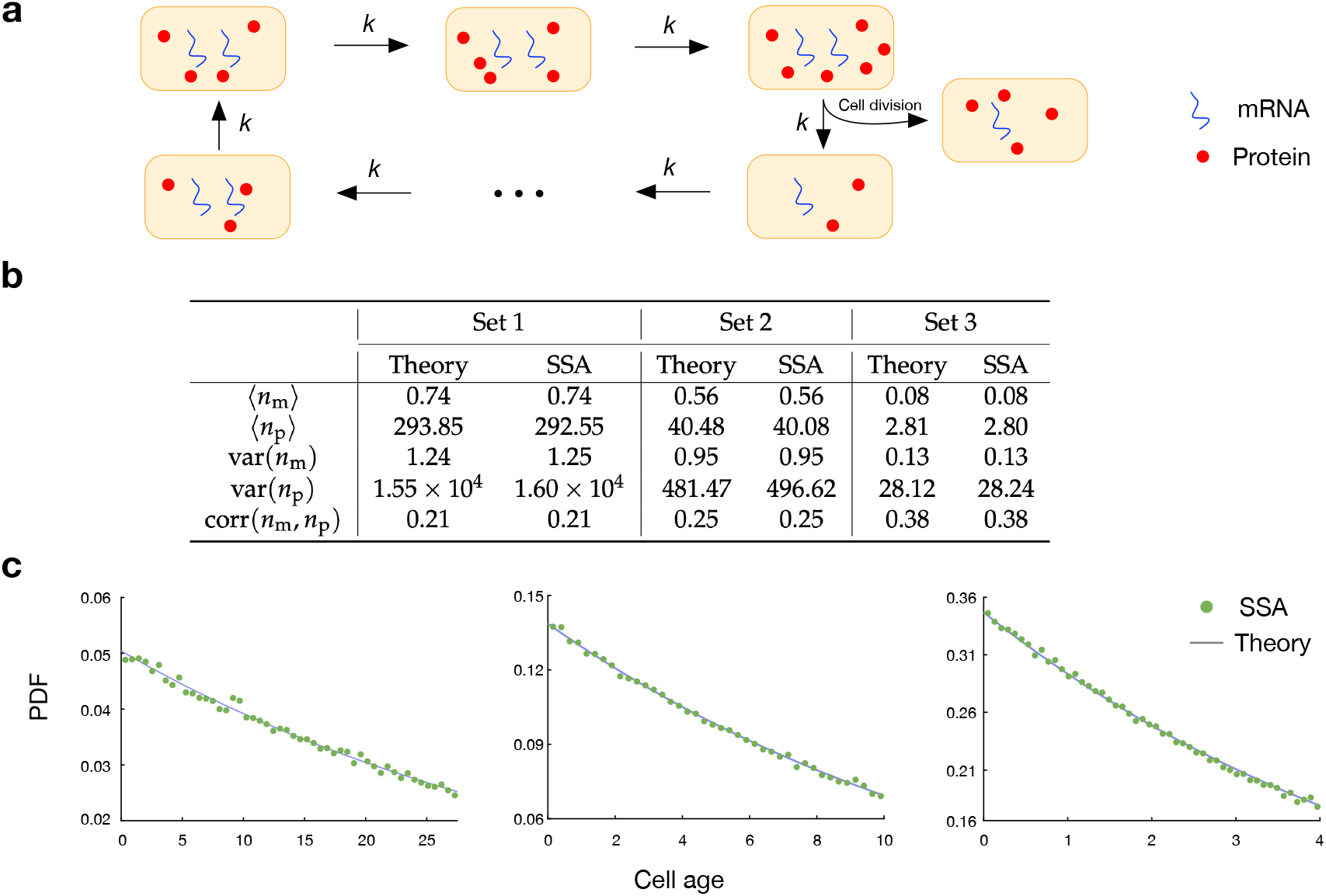
Comparison of the population snapshot predictions of Models III and IV. (a) Illustration of Model IV. This is described in detail in Appendix E. (b) The mean, variance and covariance of mRNA and protein numbers were calculated from SSA simulations of Model IV; specifically starting from a certain number of cells, the population grows exponentially and the moments are calculated after several generations when a steady-state distribution has been achieved. These moments are compared to those predicted by Model III using Eqs. (44)-(48) and excellent agreement is found between the two models for three sets of parameters. (c) From simulations of Model IV we also compute the cell-age distribution; the cell age for a cell with cell-cycle stage *i* is given by *t* = *iT/N*. The distribution from simulations is found to be in excellent agreement with the theoretical distribution *p*(*t*) = 2^1−*t/T*^ ln 2*/T* used in the analytical calculations for Model III. The kinetic parameters together with the initial number of cells *n*_initial_, the total simulation time *t*_max_, the number of phases *N*, and cell-cycle stage transition rate *k* used for SSA simulations are: (Set 1) *σ*_on_ = 0.1, *σ*_off_ = 0.1, *ρ* = 3, *λ* = 10, *d* = 2, *T* = 27.5, *N* = 50, *t*_max_ = 400, *n*_initial_ = 5, *k* = 1.82; (Set 2) *σ*_on_ = 0.5, *σ*_off_ = 2, *ρ* = 3, *λ* = 5, *d* = 1, *T* = 10, *N* = 40, *t*_max_ = 180, *n*_initial_ = 1, *k* = 4; (Set 3) *σ*_on_ = 0.1, *σ*_off_ = 2, *ρ* = 2, *λ* = 6, *d* = 1, *T* = 4, *N* = 50, *t*_max_ = 80, *n*_initial_ = 1, *k* = 12.5.

## V. SUMMARY AND DISCUSSION

In this paper, we have extended the classical three-stage model to include a cell-cycle description and solved it exactly using the generating function method. This is a theoretical advance because to the best of our knowledge, presently there is not an analytical solution for the joint distribution of mRNA and protein numbers in a three-state model of gene expression with or without a cell-cycle description. Prior to this work, an exact analytical solution for the joint distribution existed only for the classical two-stage model where there is no promoter switching and no cell-cycle description^31,32^ and the marginal distribution of protein numbers in the classical three-stage model was only derived when the timescales of certain reactions are well separated^20^.

Because our model has the added advantage of explicitly incorporating cell division whereas the classical three-state model^20^ does not, comparing the steady-state population snapshot statistics of the two enabled us to shed light on the effect that cell division has on the predictions of the Fano factor of protein number fluctuations and the correlation coefficient between mRNA and protein number fluctuations. We found there are substantial deviations between the two model predictions because the first-order reaction modelling mRNA and protein dilution in the classical model cannot capture the fluctuations induced by cell division. In particular the classical model underestimates the Fano factor of protein numbers. The same has been previously reported using a model of bursty protein production (where the burst size is sampled from a geometric distribution) and cell division^26^; however this model is a special case of Model III presented in this paper (valid only when certain parameter restrictions apply) and hence the Fano factor analysis presented here is more general. Perhaps more interestingly, we also found that the classical model can either under or overestimate the correlation between mRNA and protein number fluctuations. Roughly speaking, overestimation occurs when the ratio of the cell division time and the mRNA lifetime *d*_*m*_*T* is greater than 10 (Fig. 4c and d). This property reflects the intuitive observation that if during most of the cell-cycle the mRNA is roughly in steady-state but the protein number is mostly increasing with time (since removal occurs only at cell division), then the correlation between the two will necessarily be weak; our cell-cycle model captures this but the classical model overestimates the correlation because it predicts that both mRNA and protein reach or approach a steady-state since both decay due to a first-order reaction.

Our model of course also has limitations, the major one being the inherent assumption that protein removal only occurs via dilution and not due to degradation. It is estimated that in mammalian cells about 70% of proteins have a lifetime that is longer than the mean cell-cycle duration^35^ – for these proteins, our model is applicable because removal will be dominated by dilution. To obtain a stochastic description for the rest of the proteins, active protein degradation needs to be included but unfortunately this then makes the analytical solution of the chemical master equation challenging and it is currently unclear how to overcome this issue.

An interesting direction we have not here explored is the implication of our analytical results to parameter inference from experimental data^6,44^, i.e. given measurements of single cell mRNA and protein numbers in a population of cells^45^, how is inference of rate parameter values (by matching the theoretical and experimental joint distribution via the method of maximum likelihood) using Model III (three-stage model with cell-cycle) different than using Model I (classical three-stage model without cell-cycle)? This is under investigation and we hope to report on it in a separate study.

## VI. CODE AND DATA AVAILABILITY

The Mathematica notebooks, Python codes and relevant data are deposited at https://github.com/edwardcao3026/three-stage-model.git.

## ACKNOWLEDGMENTS

Z.C. acknowledges support from NSFC No. 62073137, Shanghai Action Plan for Technological Innovation Grant (No. 22ZR1415300, 22511104000, 23S41900500) and Shanghai Center of Biomedicine Development. R.G. acknowledges support from a Leverhulme Trust grant (RPG-2020-327).

## Appendix A: Derivation of Eq. (22)

It then follows from Eqs. (19) and (20) that the solution *G*_1_ becomes

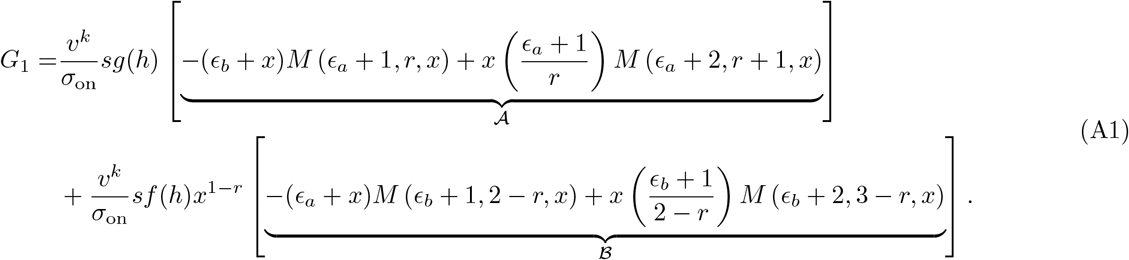

In the following steps, we show that Eq. (A1) can be further simplified by making use of recurrence relations of the Kummer function. For the term 𝒜, we have

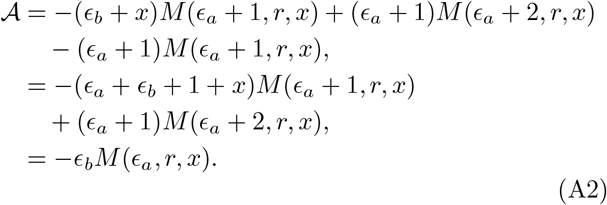

Here the first step of Eq. (A2) is achieved by applying Eq. (13.3.4) of Ref. 40 to the second term in 𝒜, and the second step re-organizes the expression in the first step, while the third step follows Eq. (13.3.1) of Ref. 40. By similar arguments, we also obtain

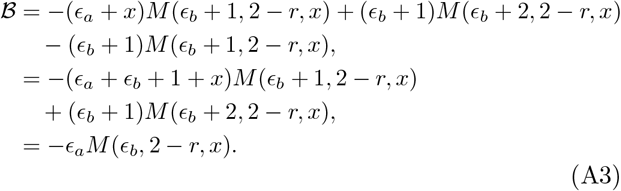

Finally using Eqs. (A1)-(A3) we obtain the much simpler expression for *G*_1_ given by Eq. (22).

## Appendix B: Derivation of Eq. (24)

The left-hand side of both equations in Eq. (23) are linear functions of both *f* and *g*, and hence they can be immediately obtained

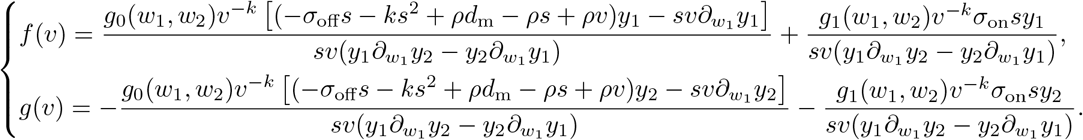

Note that the term 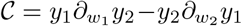 appears in several parts of the expression; it introduces the reciprocal of the Kummer function into the full solution and may thus hamper the efficient evaluation when we use *G*_0_ and *G*_1_ to obtain *P*_0_(*n*_m_, *n*_p_) and *P*_1_(*n*_m_, *n*_p_). Therefore, we need to simplify *C*, which is indeed the Wronskian identity of the two general solutions *y*_1_ and *y*_2_. Introducing 𝒟, we find 𝒞 simplifies to

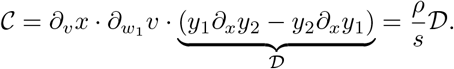

Utilizing the observations above, one can simplify *D* to

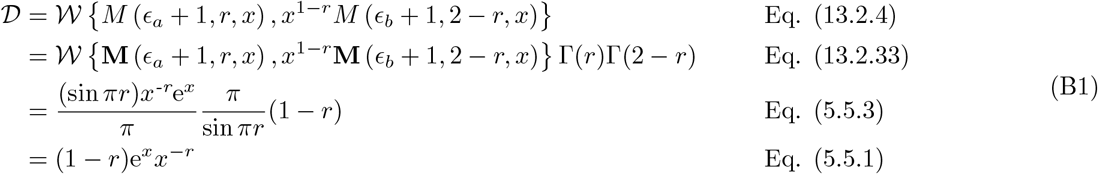

where 𝒲 {·, ·} stands for the Wronskian identity and **M**(*a, b, z*) = Γ(*b*)*M* (*a, b, z*). Note that the equations of Ref. 40 used in the above simplification steps are described at the end of each step. Following the procedures performed in Eqs. (A2) and (A3), we can use Eq. (B1) to simplify *f* and *g* to the form in Eq. (24) under the condition that *h* = *v* when *t* = 0.

## Appendix C: Marginal distribution of mRNA numbers for Model II

We start by setting *w*_2_ = 0 in Eq. (28) which leads to the simplified quantities

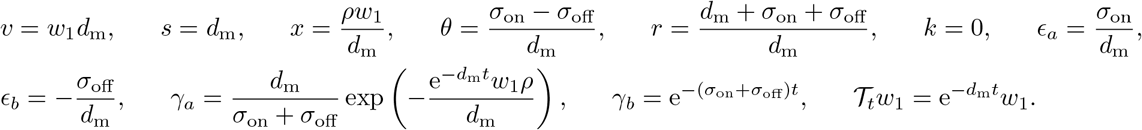

This implies the functions *k*_1_, *k*_2_, *k*_3_ and *k*_4_ in Eq. (28) have the following form

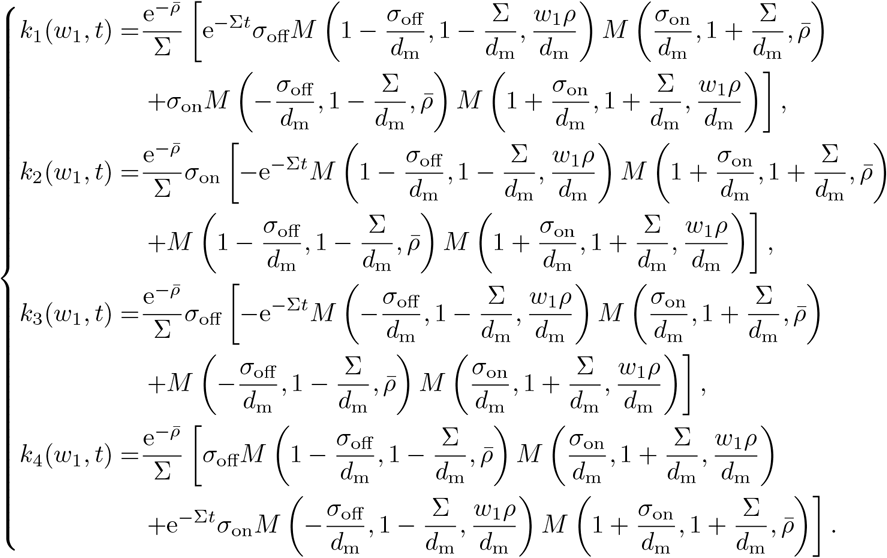

where 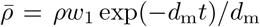and Σ = *σ*_on_ + *σ*_off_. For initial conditions of promoter state *D*_0_ and zero mRNA, specifically *g*_0_(*w*_1_) = 1 and *g*_1_(*w*_1_) = 0, one can straightforwardly verify that the solution becomes perfectly the same as Eqs. (13) and (14) in Ref. 37 (whereby *ρ*_*b*_ should be set to 0) according to *G*_0_(*w*_1_, *t*) = *k*_1_(*w*_1_, *t*) and *G*_1_(*w*_1_, *t*) = *k*_3_(*w*_1_, *t*).

## Appendix D: Marginal distribution of proteins numbers for Model II under timescale separation

We will here show the reduction of the solution Eq. (29) under the conditions *σ*_off_ ≫ *σ*_on_ and *ρ/σ*_off_ being some finite quantity. Specifically, for some finite 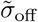 and 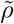, we posit the existence of a variable *δ* such that 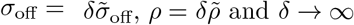 and *δ*→.∞

To achieve so, we will decompose the problem into showing the following five subproblems

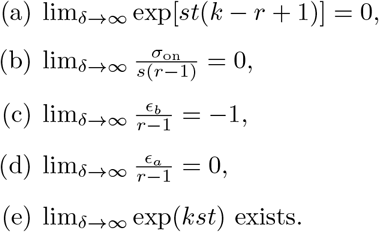

For (a), it is known that *w*_*2*_ ∈ [-1,0] since *z*_*2*_ ∈ [1,0] One immediately has

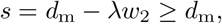

and further deduce from Eq. (14) that

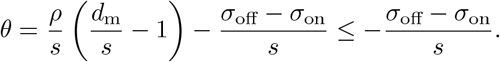

Since *σ*_off_ ≫*σ*_on_ and *σ*_off_ → ∞as *δ* → ∞, we conclude that *θ* → − ∞as *δ* → ∞. More specifically, *θ* ∼ 𝒪 (*δ*). It is also known from Eq. (14) that

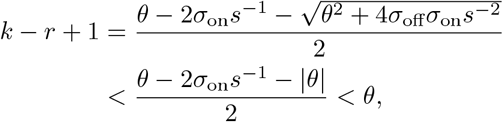

which indicates that *k* − *r* + 1 → − ∞ as *δ* → ∞. Equivalently, it means lim_*δ*_ exp[*st*(*k − r* + 1)] = 0.

For (b), since *r >* 1 and *s >* 0, we have

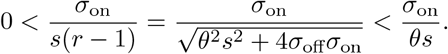

As *δ* → ∞, *σ*_on_*/*(*θs*) → 0, which suggests that

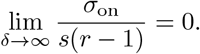

For (c), it follows from Eq. (31) that

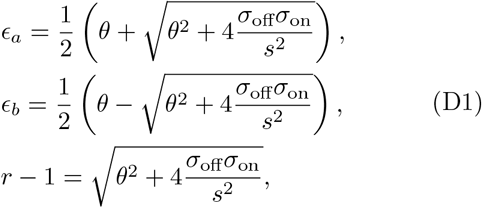

which further leads to

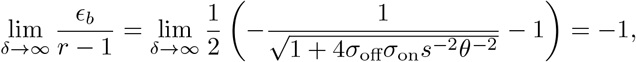

where the last step uses *θ* ∼ 𝒪 (*δ*) and *σ*_off_ ∼ 𝒪 (*δ*). For (d), it is known from Eq. (D1) that

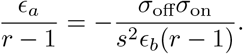

Thus, we have

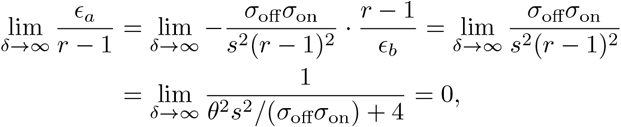

since *θ* ∼ 𝒪 (*δ*) and *σ*_off_ ∼ 𝒪 (*δ*).

For (e), we first show that *k* is a finite quantity under the condition *δ* → ∞. By using Eqs. (14) and (D1), *k* can be written as

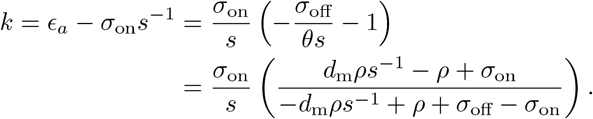

Hence, we conclude that

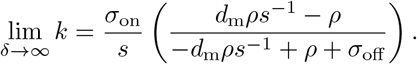

Based on the results (a)-(e), the solutions *G*_0_ and *G*_1_ reduce to

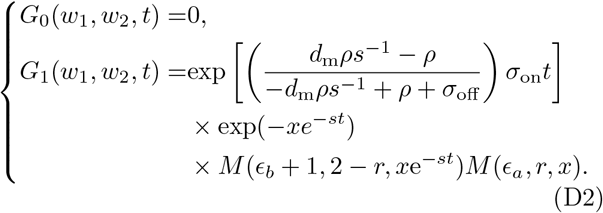

Next, we will show how Eq. (D2) can be further reduced under the condition that *d*_m_ and *λ* are very large, and *λ/d*_m_ is finite. Specifically, *d*_m_ ∼ *λ* ∼ *O*(∆) and ∆ → ∞. As such, it immediately follows that *s* ∼ *O*(∆)

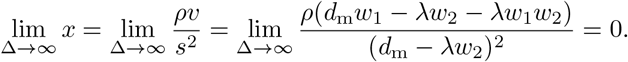

Therefore, *x*e^−*st*^ → 0 as ∆ → ∞. It is suggested by Eq. (14) that *r* is finite. And so are *ϵ*_*a*_ and *ϵ*_*b*_, according to Eq. (21). Then, by expanding Kummer functions in Eq. (D2), we obtain that

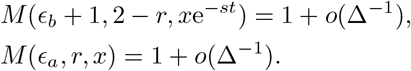

These lead to the reduced solution

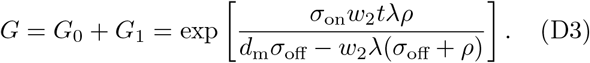

Finally, we show Eq. (D3) is the solution of the chemical master equation of the reaction system Eq. (30). To this end, we analytically solve the dynamics of this reaction system – instead of directly solving the corresponding generating function equation, we adopt a shortcut by using the generating-function property of a compound process (Supplementary Note 3 in Ref. 46).

We are interested in the distribution of protein numbers *n*_p_(*t*) to the system Eq. (30). Clearly, *n*_p_(*t*) is a random integer variable, and is determined by two factors – the number of “packages” *I*(*t*) (where a package stands for a protein-production event occurring before time *t*) and the number of proteins *X*_*i*_ Geom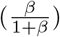 in each package *i*. Therefore, *n*_p_(*t*) can be presented in the form:

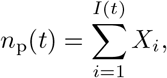

thereby constituting a compound process. The package number *I*(*t*) is determined by the pure birth process with rate *α*, and hence its generating function is

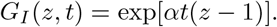

The generating function of *X*_*i*_ is

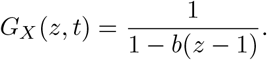

Hence, the generating function for *n*_p_(*t*) becomes

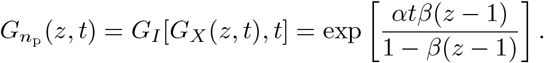

Finally substituting the expressions for *α* and *β* from Eq. (31) into the equation above, we find it leads the same equation as Eq. (D3), thereby completing the proof.

## Appendix E: Description of Model IV

Model IV is illustrated in Fig. 6a. We divide the cellcycle into *N* stages; the reactions in each stage are the same as Model II. The rate of transitioning between the stages is *k* and cell division (and binomial partitioning of gene products) occurs when the stage changes from *N* to 1. The cell-cycle duration is hence a sum of *N* exponential variables with mean 1*/k*, i.e. the cell-cycle duration is Erlang distributed with mean *T* and coefficient of variation 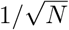. Note that the cell-cycle stages are not to be understood as the actual biological cell-cycle phases (*G*_1_, *S, G*_2_, *M*); rather these are introduced as a natural means to model an Erlang distributed cell-cycle duration where the parameter *N* can be directly estimated from experimental measurements of the cell-cycle duration distribution^27^.

Simulations of this model starting from one or more cells leads to an exponentially growing population of cells. After some generations the system reaches a steady-state, in the sense that the population distribution of gene products becomes invariant with time. From a snapshot measurement, we compute the moments of gene products which we compare with those of Model III in Fig. 6b. The cell-age distribution is also computed and it is found to be in good agreement with the cell-age distribution used in calculations of Model III (Fig. 6c).

